# Genetic characterization of a captive marmoset colony using genotype-by-sequencing

**DOI:** 10.1101/2023.06.22.545969

**Authors:** SA Cole, MM Lyke, C Christensen, D Newman, A Bagwell, S Galindo, J Glenn, D Layne-Colon, K Sayers, SD Tardif, LA Cox, CN Ross, IH Cheeseman

## Abstract

The marmoset is a fundamental non-human primate model for the study of aging, neurobiology, and many other topics. Genetic management of captive marmoset colonies is complicated by frequent chimerism in the blood and other tissues, a lack of tools to enable cost-effective, genome-wide interrogation of variation, and historic mergers and migrations of animals between colonies. We implemented genotype-by-sequencing (GBS) of hair follicle derived DNA (a minimally chimeric DNA source) of 82 marmosets housed at the Southwest National Primate Research Center (SNPRC). Our primary goals were the genetic characterization of our marmoset population for pedigree verification and colony management and to inform the scientific community of the functional genetic makeup of this valuable resource. We used the GBS data to reconstruct the genetic legacy of recent mergers between colonies, to identify genetically related animals whose relationships were previously unknown due to incomplete pedigree information, and to show that animals in the SNPRC colony appear to exhibit low levels of inbreeding. Of the >99,000 single-nucleotide variants (SNVs) that we characterized, >9,800 are located within gene regions known to harbor pathogenic variants of clinical significance in humans. Overall, we show the combination of low-resolution (sparse) genotyping using hair follicle DNA is a powerful strategy for the genetic management of captive marmoset colonies and for identifying potential SNVs for the development of biomedical research models.

## INTRODUCTION

The common marmoset *(Callithrix jacchus)* is an important non-human primate (NHP) model for biomedical research (Miller, 2017; Ross & Salmon, 2019; Servick, 2018). The marmoset provides unique practical advantages relative to other NHPs, including small body size, rapid reproductive maturation, high fecundity, compressed life cycle, and comparative ease of handling (Abbott et al., 2003; Kishi et al., 2014; Tardif & Ross, 2019; Tardif et al., 2003). Additionally, aspects of their social behaviors and communication closely resemble those observed in humans (Miller et al., 2016). These advantages have contributed to the recent use of marmosets for developing new disease models using gene editing techniques and translational studies focused on gene therapy (Kishi et al., 2014; National Academies of Sciences & Medicine, 2019). Marmosets have also been utilized as a model species for a number of research areas, including aging, neuroscience, and infectious disease (National Academies of Sciences & Medicine, 2019), and new applications continue to be explored.

Given this increasing importance, there has been recent interest in genetic characterization of existing captive colonies to develop colony management strategies that maintain their genetic diversity and long-term value (reviewed in Harding 2017). Several approaches have been used for population genetic characterization of NHPs, including whole genome sequencing (WGS), whole exome sequencing (WES), and genotype-by-sequencing (GBS) (Bimber et al., 2016). The first whole genome sequence arising from a common marmoset was generated from the genome of an animal housed at the Southwest National Primate Research Center (SNPRC) (MGSAC, 2014). The genome was sequenced using Sanger sequencing (6x) and a whole-genome shotgun approach (MGSAC, 2014). The marmoset genome assembly was then improved through deep sequencing of a marmoset housed in Japan using next generation sequencing (NGS) (Sato et al., 2015), and subsequent efforts continue to increase the quality and coverage (Rogers & del Rosario, 2019).

An important aspect of the genetic characterization of NHPs as models for human health and disease is the identification of functional single-nucleotide variants (SNVs) (Cline & Karchin, 2011; Xue et al., 2016). SNVs represent common genetic variation that can be useful for identifying variants that influence gene functions (Jasinska, 2020). Of particular interest are SNVs located in genes with known phenotypic consequences associated with health and disease in humans. Potentially functional SNVs have been identified and cataloged for several NHP species, including rhesus macaques (*Macaca mulatta*; Bimber et al. 2017, Xue et al. 2016) and vervet monkeys (*Chlorocebus sabaeus*; Huang et al. 2015) with the goal of identifying sites relevant to studies of human health, though this has not yet been extensively done for marmosets. Once identified, these variants can be explored as potential orthologs to human disease genes, further facilitating the development of effective marmoset models for biomedical research.

A special consideration when interpreting marmoset genomic data is that marmoset littermates are chimeric (Benirschke et al., 1962). Marmoset tissues have a range of chimerism, with hematopoietic-derived tissues showing more extensive chimerism (Ross et al., 2007; Sweeney et al., 2012; Takabayashi & Katoh, 2015). To minimize the effect of chimerism when assessing genetic diversity based upon sequence analyses (sequencing or genotyping), investigators choose tissues with minimal chimerism such as hair, skin, or nail (Silva et al., 2017; Takabayashi & Katoh, 2015). Due to the low yield of DNA derived from some of these sources, cultured fibroblasts have been used for more comprehensive molecular approaches such as WGS (NPRC Marmoset Genomics Working Group). To minimize chimerism in our samples, we extracted DNA from hair follicles, which have far lower levels of chimerism than blood (Ross et al., 2007) and yield more DNA of higher quality than finger nails.

The marmoset colony housed at the SNPRC is particularly valuable because of its genetic history. It is an outbred population that can be traced for up to 12 generations, with a founder population of 120 and a current population of ≈400. Based upon pedigree, the SNPRC population is particularly diverse, with founders from the University of Zurich, the University of London, Marmoset Research Center-Oak Ridge (MARCOR), Osage Research Primates (ORP), Harlan, Wisconsin National Primate Research Center (WNPRC), NIHCHD (National Institutes of Health-Child Health and Development), New England NPRC (NEPRC), and Worldwide Primates (WWP). The SNPRC colony manager uses pedigree analyses and relatedness tools to evaluate and create breeding pairs that protect the genetic diversity of the colony. The current research will increase our knowledge of the genetic diversity and relatedness of these important primate resources.

The goal of this study was to develop a genome-wide genetic resource for the SNPRC pedigreed marmoset colony that can be used in a cost-effective manner for: (1) pedigree verification; (2) assisting with colony management; and (3) informing the scientific community regarding the functional genetic makeup of this valuable colony resource. Although WGS costs continue to decrease and low coverage WGS can be done relatively inexpensively, confidently calling variants in potentially chimeric samples requires at least moderate coverage. For initial development of this resource, we thought it was important to provide high resolution single nucleotide variant data in exons, which are typically the highest priority SNVs for investigators. While WES shows promise in characterizing NHP genomic variation (Chan et al., 2021) commercially available WES currently uses human DNA sequences for exon capture, and exons for some marmoset genes would be missed. For example, Chan and colleagues (2021) showed that 6.5% of the coding exons, 32% of the 5’ UTR exons, and 9.8% of the 3’ UTR exons were not captured using the human exon capture kit. Therefore, we chose to use a species agnostic GBS approach where we can enrich for coding regions of the genome by restriction enzyme selection and generate moderate coverage for each read at about one-fifth the cost per sample of WES.

## METHODS

### SNPRC marmoset colony

The SNPRC marmoset breeding colony was originally established in 1993 by Dr. Suzette Tardif at the University of Tennessee-Knoxville with ten founding animals imported from Zurich, Switzerland (University of Zurich) and 35 from the United Kingdom (UK; University of London). The colony added animals from MARCOR, ORP, and the NICHD colony prior to moving the colony to the SNPRC in 2001. Additional animals were imported for projects and production starting in 2005 including WNPRC, Harlan, NEPRC, NIH and WWP resulting in 120 founding animals. In 2015 the colony was further expanded through a merger between the SNPRC and NEPRC colonies, introducing more than 180 new animals to the SNPRC colony. Several marmosets have also returned to the SNPRC colony following sales to outside institutions. Most of the marmoset colony belongs to one pedigree, which includes >3,000 animals (≈400 living) spanning 12 generations in depth as of April 2023. Hair follicle DNA was collected from 82 animals for GBS, and used for sequencing and evaluation. This research protocol was approved by the Texas Biomedical Research Institutional Animal Care and Use Committee #927CJ.

### Isolation of DNA

For this study, hair follicle samples were opportunistically collected from 82 living animals. Hair and follicles were collected at the time of a physical exam or of euthanasia. A site on the body was chosen that was visually free of scent marking secretions, such as the tail or lower back. A clump of about 50-100 hairs was grasped with clean forceps and pulled to remove hair with the follicles intact. The clump was placed follicle end first into a sterile screw cap tube and frozen at - 80C. For DNA isolation, the hair and follicles were rinsed in 2mls of PBS (2x) prior to the start of the DNA isolation procedure, which used the QIAmp DNA Mini Kit (Qiagen) following the manufacturer’s protocol for tissues, including the optional RNase step.

### Genotype-by-sequencing (GBS)

GBS was completed using the DNA from 82 individual marmosets. GBS libraries were prepared following the GBS method developed by Elshire et al. (2011) and optimized for the marmoset genome. In brief, 50 ng of genomic DNA was digested in 20 µl reactions with 4U *Ava*II. Adapters with barcodes from 4-9 bp in length, designed using the GBS Barcode Generator (deenabio.com/services/gbs-adapters) were then ligated to the digestion products in 50 µl reactions using 400 cohesive end units of T4 DNA ligase (NEB). Following ligation, samples were pooled and purified using the QIAquick PCR Purification Kit (Qiagen). Pooled DNA libraries were amplified in 50 µl reactions with 1x NEBNext High Fidelity PCR Master Mix (NEB) and 12.5 pmol PCR primers containing complementary sequences for adapter-ligated DNA. Temperature cycling consisted of 72° for 5 min, 98° for 30 s, followed by 18 cycles of 98° for 10 s, 65° for 30 s, and 72° for 30 s, with a final extension at 72° for 5 min. Amplified libraries were purified using the QIAquick PCR Purification Kit (Qiagen) and the quality and quantity of each library assessed using the Agilent DNA 1000 chip (Agilent Technologies) and KAPA Library Quantification Kit (Kapa Biosystems), respectively. The final DNA libraries were hybridized to Rapid Run Flow Cells (Illumina) for cluster generation using the TruSeq™ PE Cluster Kit (Illumina) and sequenced with the TruSeq™ SBS Kit (Illumina) on an HiSeq 2500 (Illumina) using a 150-cycle paired-end sequencing run.

Sequence reads were demultiplexed with GBSX v1.0.1 (Herten et al. 2015; github.com/GenomicsCoreLeuven/GBSX). Sequence data were imported into Partek Flow (Partek, Inc.). Sequence reads were trimmed and filtered based on a minimum quality score (Phred) of 30 and a minimum read length of 25. The filtered reads were aligned to the marmoset C_jacchus3.2.1 assembly using BWA-MEM alignment tool (Li, 2013) Single nucleotide variants (SNVs) were detected using a minimum Phred quality score of 30 and filtered further using GATK to include only those that had a minimum read depth of 5 and a minimum log-odds ratio of 3.05 to ensure high-quality variants. SNVs were annotated and effects predicted using Ensembl Variant Effect Predictor (VEP, McLaren et al. 2016; useast.ensembl.org/info/docs/tools/vep/ index.html). Sequences have been deposited in NCBI with Accession numbers SRR18101012-SRR18100931.

### Generating a high confidence genetic database

We retained autosomal, biallelic SNV data from the hair follicle DNA samples, and loaded the data into SOLAR (Almasy and Blangero 1998) for initial evaluation of allele frequencies and descriptive tallies. A modified pedigree based on the 82 original samples, their parents, and their descendants was developed for use with all pedigree-based analyses (n=822). One of these animals was unrelated to the others and was therefore not included in pedigree and kinship estimation, though included in all other analyses. INFER, which is a program within the PEDSYS software system (Dyke, 1996), examines each offspring-father-mother triplet and, when possible, adds missing alleles and genotypes according to the Mendelian laws of transmission. The program iterates through the pedigree as many times as necessary until no more assignments can be made. The inferred data were then combined with the pedigree data for further analyses. Following the inference of new genotypes, we used SimWalk2 (Sobel et al., 2001) to identify and remove genotypes inconsistent with Mendelian properties within families including the grand-parental generation, as well as distributions of alleles within entire sibships. SimWalk2 reports the *overall* probability of mistyping at each observed genotype (in fact, at each observed allele). When genotypes were flagged with a significant probability of mistyping, they were removed from the dataset. The resultant data file was again evaluated with SOLAR (Almasy & Blangero, 1998) to create a summary list of all variant loci. This summary provided the number of samples counted per variant, the SNV major and minor allele frequencies and the associated p-values of a test of whether the Hardy-Weinberg equilibrium (HWE) holds. From the summary list of all variant loci, we selected and removed all SNVs that had low call rates keeping only those SNVs where 95% of samples (78 of initial 82) were typed. The remaining set of annotated variants represent the high confidence database carried forward for further analyses.

### Statistical and Genetic Analysis

We performed inference of historical admixture in the colony with ADMIXTURE v1.3.0 (Alexander et al., 2015) using high quality SNVs from the 82 animals directly genotyped in this study. In addition, we have substantial genotype data from 48 animals where genotypes were inferred by inheritance in INFER, which were also included to increase coverage of the pedigree. ADMIXTURE analysis was run unsupervised on these 130 animals using 1-5 theoretical ancestral populations (*K*). Values of *K*>3 failed cross validation and were discarded. We then integrated publicly available WGS data from 9 animals from WNPRC (*n*=2), NEPRC (*n*=2), and SNPRC (*n*=5) (MGSAC, 2014). We ran a supervised analysis (*K*=3) using the WGS samples incorporating the population labels from each primate center.

We estimated kinship from genetic data using lcMLkin (Lipatov et al., 2015), using high quality genotypes from 81 animals with GBS data (excluding the individual that was unrelated to the rest of the pedigree). lcMLkin estimates kinship while accounting for incomplete data and the reduced ability to capture both alleles at a given locus. Additionally, empirical kinship was directly inferred from the pedigree records of 3,232 animals using the kinship2 package in R v4.2.2 (Sinnwell et al., 2014). We found a strong correlation between mean read depth and the proportion of heterozygous base calls. To account for this dependency in our analysis we performed linear regression of proportion of heterozygous base calls using mean read depth and generation as covariates. This was implemented in the lm function in R.

To investigate functionally significant genetic variation, we assessed the overlap of the GBS SNVs with gene regions associated with immune function and inflammation, neurological traits, aging, obesity, and diabetes using the NCBI “Gene” database (https://www.ncbi.nlm.nih.gov/gene). The NCBI Gene database contains the known functions of genes of many different species, including humans and NHPs. We also merged our GBS-identified SNVs with the human ClinVar database (ncbi.nim.nih.gov/clinvar) to identify SNVs that were located in genes with identified human variants that result in phenotypes with potentially clinical outcomes.

## RESULTS

### Genotype-by-sequencing identifies SNV with potential functional significance in marmosets

Forty-five percent of reads aligned to the transcriptome. As the transcriptome is 3.3% of the marmoset genome, this indicates that the choice of restriction enzyme to target coding regions was appropriate for generating gene-centric GBS data. The GBS data from the 82 marmoset hair follicle DNA samples yielded 231,317 biallelic SNV loci. Excluding loci from chromosomes X and Y yielded 216,015 SNVs. After implementing our quality control procedures which include Mendelian error cleaning and including only SNV with high call rates, we obtained a high quality set of SNV for further analysis. Table 1 lists the types of potential functional effects identified for this marmoset SNV dataset. We assessed the overlap of the GBS SNVs with gene regions associated with immune function and inflammation, neurological traits, aging, obesity, and diabetes in humans using the NCBI Gene database. We identified 7,738 SNVs associated with immunity/inflammation, 289 associated with neurological traits, 2,544 with aging, 5,715 with obesity, and 5,402 with diabetes. We also merged our GBS-identified SNV with the human ClinVar database (ncbi.nim.nih.gov/clinvar) and identified 9,897 SNVs that were located in genes with identified human variants that result in phenotypes with potentially clinical outcomes (Table 1).

**Table 1.**
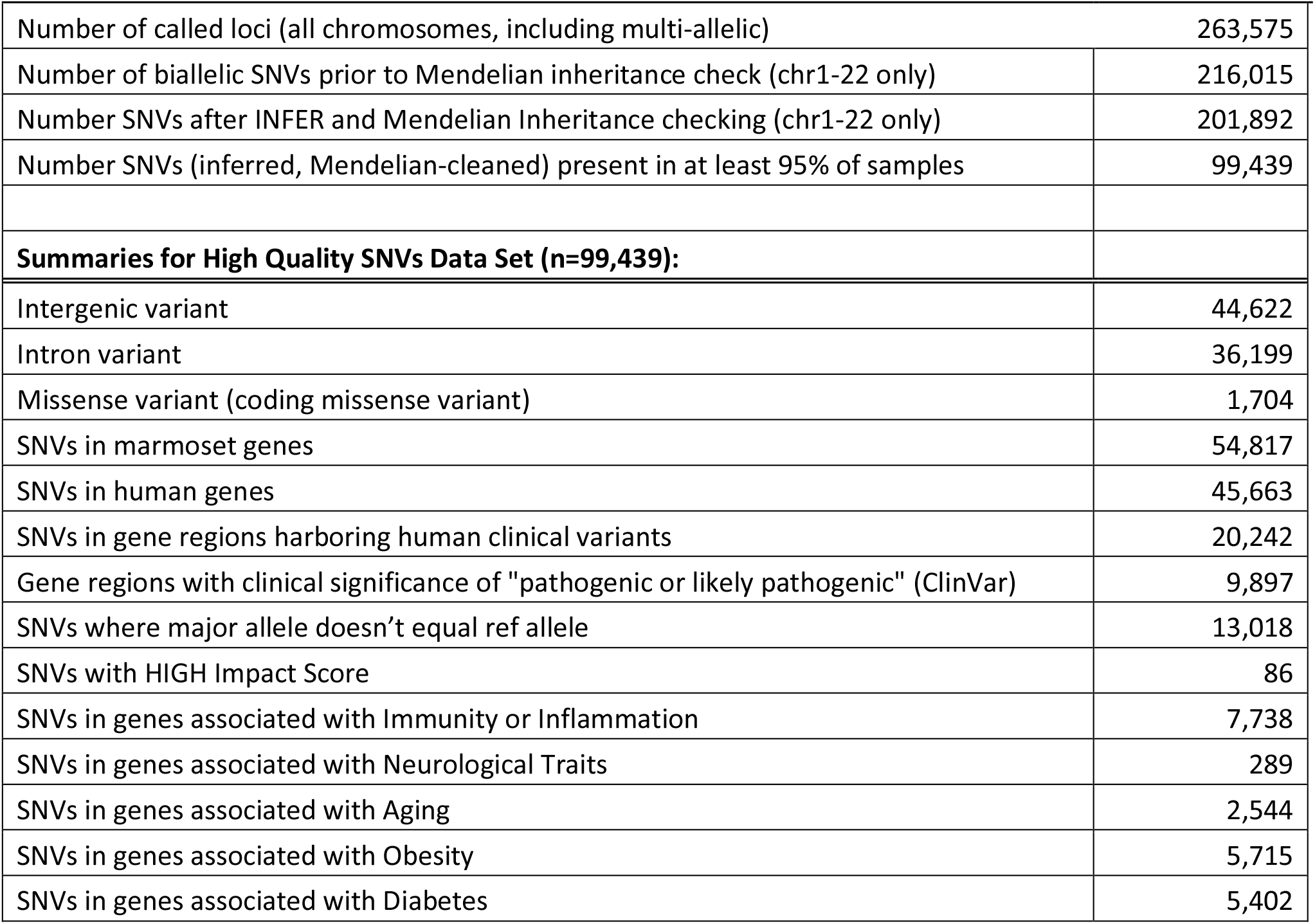
Genome-wide GBS data from common marmosets (*Callithrix jacchus*) housed at the SNPRC.

### Admixture analyses reveal the impact of past mergers on colony structure

To assess the impact of the colony mergers, we combined cleaned, imputed GBS data from the 82 animals sequenced here with previously reported whole genome sequencing data and imputed genotypes inferred solely from pedigree structure. For admixture analyses, we lowered our threshold for SNV filtering to include 131,648 biallelic SNVs typed in >50% of individuals. We estimated the number of ancestral populations giving rise to genetic diversity in the SNPRC colony and the prevalence of ancestry components in the joint set (GBS and inferred) of 130 animals using ADMIXTURE. Cross-validation of cluster numbers *K*>3 did not converge and were excluded. At *K*=2 (the best supported number of clusters), SNPRC animals showed a clear bimodal distribution of ancestry with 60.8% of animals deriving >95% of their ancestry from a single component (Fig. 1A). These results were also supported at *K*=3 (Fig. 1B). Results of the supervised admixture analysis with population labels for each primate center also indicate that the ancestry of most animals is derived from a single population (Fig. 1C). Figure 1D shows the known pedigree with each animal included in the admixture analysis shaded by the proportion of NEPRC ancestry. There is a clear subdivision in the pedigree which recapitulates the colony merging. Our data allows us to assess admixture in the SNPRC colony and capture the impact of past colony mergers on population structure.

**Figure 1.**
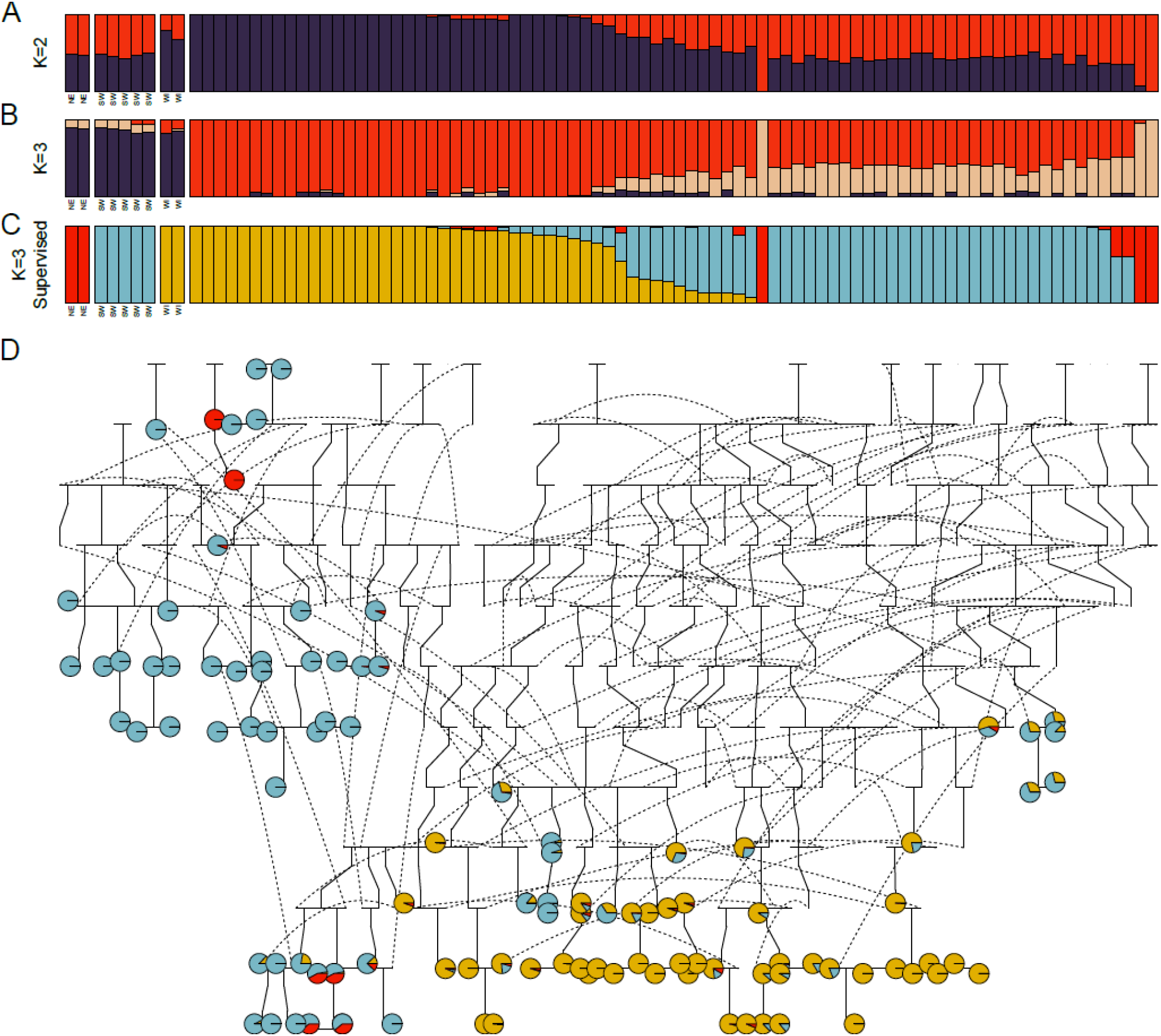
Admixture in the SNPRC colony. High quality SNVs from GBS from 82 marmosets were integrated with whole genome sequencing from NEPRC (columns 1-2, n=2), SNPRC (columns 3-7, n=5) and WNPRC (columns 8-9, n=2) to infer recent ancestry of pedigreed animals in the USA. Each bar shows the inferred ancestry proportions of each animal specifying either Fig 1A, 2 ancestral populations (K=2, top panel and best cross-validation score) or Fig 1B, 3 (K=3, middle panel). Cross validation for K>3 ancestral populations failed. In Fig 1C, ancestry was independently inferred in a supervised analysis using the population labels shown below the first 9 bars and corresponding to each NPRC. Each analysis showed a subdivided population, where most animals have a major ancestry component from a single population. Fig 1D shows the known pedigree with each animal included in the ADMIXTURE analysis shaded to represent the proportion of NEPRC, SNPRC, and WNPRC ancestry. Dashed lines show where an individual is placed multiple times in the figure. Given the complex nature of large pedigrees this is an artifact of plotting data.

### GBS data augments colony pedigree data

For colony management, genetic data directly captures inbreeding and relatedness, and can be used to inform breeding strategies and correct or enhance pedigree records. We estimated kinship directly from the genetic data for the 82 animals with GBS data using lcMLkin. We found this to have strong correlation to pedigree records (*r*^*2*^=0.78, *p*<2.2×10^-16^, Fig. 2), with a moderate global inflation of kinship estimate (intercept from a linear model=0.036, Fig. 2A). We identified 12 animals who showed no observed kinship (<0.01) from pedigree records, though were close relatives from genetic data 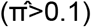. Subsequent inspection of the pedigree and animal transfer records revealed the relationships of 11 of these animals (Fig. 2B). We generated a heat map to compare the pedigree and GBS based estimates of relatedness (Fig. 2C). While both estimates capture close familial relationships, GBS based estimates of relatedness show generally higher estimates of relatedness (Fig. 2C).

**Figure 2.**
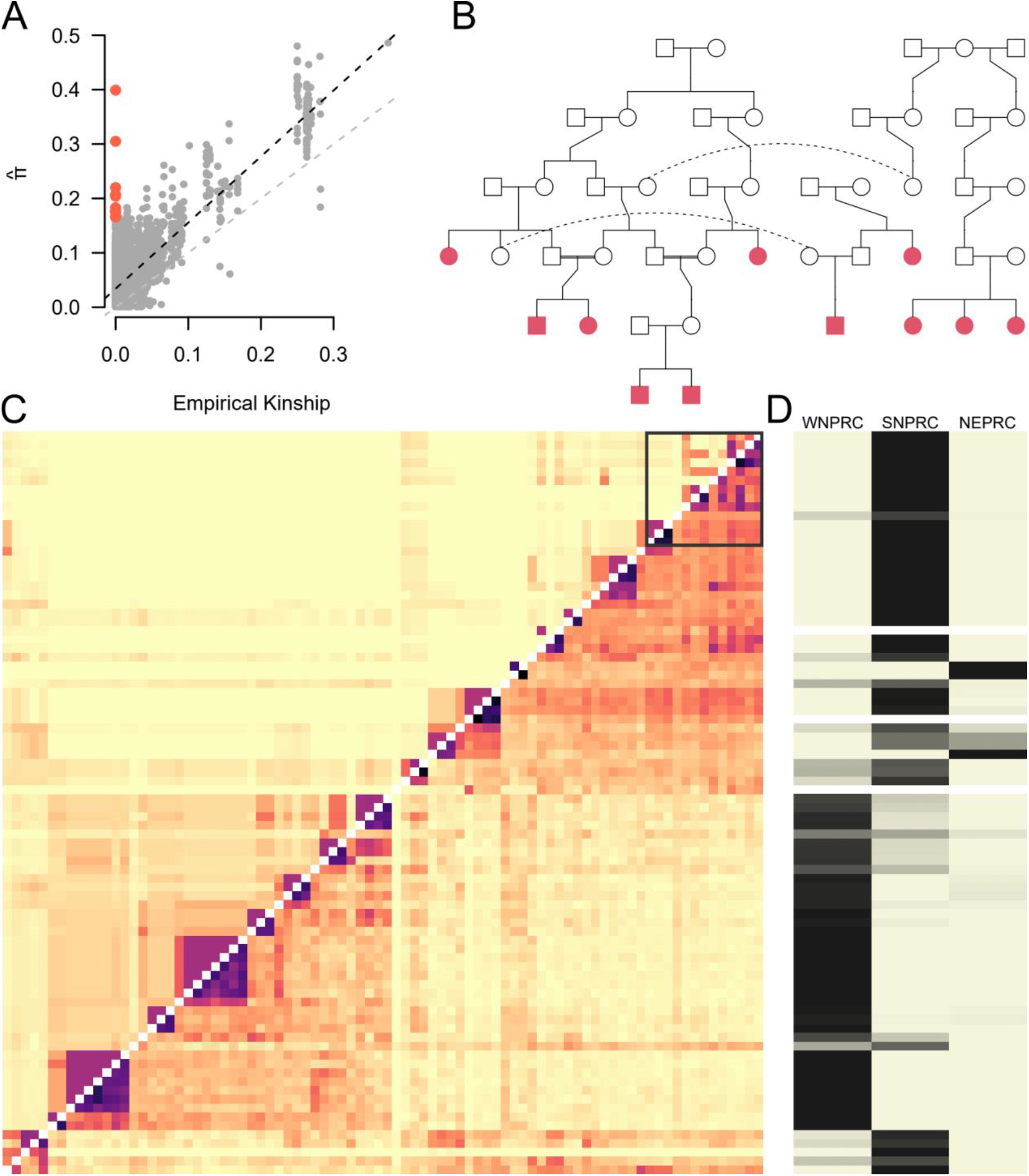
Inference of relatedness from GBS data. (A) a comparison of empirical kinship (x axis) and relatedness inferred from GBS (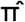, y axis). The grey dashed line shows the expectation from a perfect correlation between approaches, the black dashed line shows the observed relationship between approaches. Highlighted in the red dots are comparisons between 12 individuals which are discordant between approaches. (B) Pedigree of 11 of the 12 individuals uncovered using GBS (C) A heatmap showing the pedigree (upper triangle) and GBS (lower triangle) based estimates of relatedness. Darker colors denote a higher degree of relatedness between individuals. Highlighted by a box in the top right corner are the individuals shown in red in (A). The large difference in estimates of relatedness between the GBS and pedigree based estimates is driven by incomplete pedigree information from a single individual subject to migration between colonies. (D) Inferred ancestry components for each individual, darker colors represent the proportion of ancestry from Fig 1C.

### Genetic diversity is not declining in the SNPRC colony

A major concern in captive pedigrees is the erosion of diversity over time due to inbreeding. We tested if there was a decline in the proportion of heterozygotes in successive generations in our data. As there is a significant concern that read depth will influence the ability to accurately identify heterozygous sites, we fit a linear model on the percentage of heterozygous sites in a sample against the mean read depth and the pedigree generation. Both read depth and pedigree generation were significant predictors of percentage of heterozygous sites (Table 2). Notably, a more complex model including ancestry components did not show ancestry to be a significant predictor suggesting the proportion of heterozygous sites was not variable between founding populations. In both cases (increased read depth and more contemporary generations) there was a positive relationship with percentage of heterozygous sites. While we do not explicitly derive an estimate of the heterozygosity here, the average proportion of heterozygous sites in this population is not decreasing, suggesting this is a genetically healthy breeding population minimally impacted by inbreeding.

**Table 2.** Linear regression of proportion of heterozygous sites against pedigree generation and mean depth.

|  | Estimate (std error) | *t*-value | *p*-value |
| --- | --- | --- | --- |
| Generation | 0.03 (0.007) | 4.708 | 1.05x10<sup>-5</sup> |
| Mean Depth | 1.95 (0.16) | 12.297 | 4.92x10<sup>-20</sup> |

## DISCUSSION

Marmosets have become important resources for biomedical research (Marini et al., 2018). They have been used as models for studying a range of human conditions, from aging to infectious disease (Miller, 2017; Ross & Salmon, 2019). Marmosets are phylogenetically more distant from humans than catarrhine primates, but there are several traits that make them better suited for specific types of research. Their relatively shorter lifespans make them ideal models for studies of primate aging (Ross & Salmon, 2019). Their higher fecundity and tendency towards twinning are useful for studying aspects of pregnancy and for maintaining colony numbers (Abbott et al., 2003; Tardif & Ross, 2019; Tardif et al., 2003). Additionally, their small body size and relative ease of handling make housing and caring for marmosets more practical than other NHPs (Abbott et al., 2003; Miller, 2017). The marmosets at the SNPRC comprise a controlled breeding population with multi-generational pedigree data, making them ideal subjects for genetic characterization. With the recent increase in genome sequencing of NHPs commonly used in biomedical applications, the identification of SNVs corresponding to genes with known human health outcomes has been highly beneficial. Using GBS on hair follicle DNA, we identified 216,015 high quality SNVs in 82 marmosets from the SNPRC colony (Table 1). After quality control including Mendelian error correction and including only SNV with a high call rate and further filtering using public databases, we were able to identify variants associated with immune function and inflammation (n=7,738), neurological traits (n=289), aging (n=2,544), obesity (n=5,715), and diabetes (n=5,402) in humans (Table 1). Identifying potentially functional health related SNVs allows researchers to focus on specific regions of the genome and model genetic mechanisms of human disease.

One of the key factors when considering animals as subjects for translational research is the overall genetic health of the population (Harding, 2017; Haus et al., 2014). It is important to minimize potential adverse health effects related to inbreeding that could confound experimental design and results (Haus et al., 2014; Honess et al., 2010). Inbreeding and decreased genetic diversity are concerns when breeding any captive population as gene flow is generally limited (Harding, 2017; Haus et al., 2014). Early genetic studies suggested that overall genetic diversity of marmosets and other callitrichids was historically low (Dixson et al., 1988; Faulkes et al., 2003; Forman et al., 1986; Nievergelt et al., 2000; Watkins et al., 1991), so developing a low cost yet effective way to measure and track genetic diversity in our population has been a priority. Using the GBS data and the proportion of heterozygous sites as a measure of diversity, here we have shown that the individuals in the primary pedigree at SNPRC are genetically healthy and in general the colony appears to exhibit low levels of inbreeding. This is likely due to the diverse provenances of our founding population and the ongoing efforts of our colony manager and others to reduce the loss of diversity. As shown in our data, both read depth and pedigree generation were predictors of the percentage of heterozygous sites, and both showed positive relationships (Table 2). Our GBS approach and data QC ensured that we had a high confidence set of SNVs for analyses, and caution that low pass sequencing methods may not reveal all informative genetic variation. Regarding the effect of pedigree generation on heterozygosity, this was likely due in part to recent colony mergers that introduced new individuals into the population. In general, these migrations increase genetic diversity and decrease inbreeding, as was the case with the marmoset colony mergers discussed above.

While migrations and animal transfers can help maintain or increase genetic diversity, when animals are moved between NPRCs or other organizations, potential relationships between individuals can be missing from the pedigree record. When new animals are imported, they are considered to be unrelated to others in the colony, as there is no known relationship. Our GBS data uncovered previously unknown relationships in twelve individuals in our colony (Figure 2). Subsequent inspection of pedigree records allowed us to uncover some of those relationships. For example, a male marmoset with several offspring in the SNPRC colony was transferred to another location, where he then had offspring. One of his male offspring (CJTXGBS00082) was transferred back into the SNPRC colony, where he had unknown half-siblings. Additionally, one or both parents of three of the individuals (CJTXGBS00060, CJTXGBS00062, CJTXGBS00066) were part of the founding population, and their relationships to each other were only revealed with the GBS data. Prior to generating the GBS data, estimations of kinship and relatedness were based solely on pedigree, which is extensive and is a valuable tool, but with some inherent limitations due to potentially missing data. We have demonstrated that genetic data can play a pivotal role in identifying potential errors in the pedigree for further exploration. These related individuals, or their close relatives, might otherwise have been picked as breeding pairs, inadvertently increasing levels of inbreeding in the population.

Another outcome of importing animals from other colonies is that genetic admixture occurs. Geographically isolated populations tend to have genetic signatures due to changes in allele frequencies based on local adaptations (Cheng et al., 2022). Genetic signatures can be developed over long evolutionary periods and detected at the species level, but they can also be driven by short term population isolation, such as in captive breeding colonies (MGSAC, 2014; Schoener, 2011). When discrete populations are brought into contact and interbreed, genetic admixture occurs. The extent of admixture can be measured over subsequent generations using genetic data, such as GBS (Alexander et al., 2015). The ability to track admixture and ancestry is beneficial when analyzing potential variation in disease risk and response when developing animal models for biomedical research (Shriner, 2017). Here we were able to successfully identify genetic signatures corresponding to colony of origin using admixture analyses of our GBS data. We ran both supervised and unsupervised analyses and both reflected population structure at the colony level. Because of their relatively short evolutionary histories and rapidly shifting allele frequencies, migration of animals among NPRCs is driving population structure.

An important component of NHP colony management is the assessment and maintenance of genetic health and diversity. This is especially critical for animals used in biomedical research, where certain genetic traits may influence disease susceptibility or outcomes (Haus et al., 2014; Honess et al., 2010). It is also critical to maintain genetic diversity for the endurance and expansion of the colony itself as a long-term research resource. For marmosets, the choice of tissue used for genetic analyses requires special consideration due to their chimeric nature. Hair follicles are among the least chimeric tissues, are collected non-invasively, and yield high quality DNA suitable for advanced genetic sequencing (Ross et al., 2007). Recent years have seen an increase in commercially available sequencing options with decreasing costs, yet some methods, such as WGS, are still cost prohibitive for population-level genetic screening, especially for those colonies with limited resources. We have demonstrated that GBS provides high quality, affordable data using sparse genetic characterization for population management and for assessing ancestry and colony genetic health. The combination of hair follicle DNA with GBS represents a successful, non-invasive, cost-effective approach for colony management and for understanding evolutionary diversity of captive marmosets being used in biomedical applications.

## Financial Support

This research was supported by NIH-NCRR grant P51 RR013986 to the Southwest National Primate Research Center.

## Conflict of interest

The authors have declared no conflict of interest.

## Notes

### Competing Interest Statement

The authors have declared no competing interest.

